# Uncoupling differential water usage from drought resistance in a dwarf Arabidopsis mutant

**DOI:** 10.1101/2021.11.25.470014

**Authors:** Daniel N. Ginzburg, Flavia Bossi, Seung Y. Rhee

**Author notes:** Corresponding author: Seung Y. Rhee.

## Abstract

Understanding the molecular and physiological mechanisms of how plants respond to drought is paramount to breeding more drought resistant crops. Certain mutations or allelic variations result in plants with altered water-use requirements. To correctly identify genetic differences which confer a drought phenotype, plants with different genotypes must be subjected to equal levels of drought stress. Many reports of advantageous mutations conferring drought resistance do not control for soil water content variations across genotypes and may therefore need to be re-examined. Here, we reassessed the drought phenotype of the *Arabidopsis thaliana* dwarf mutant, *chiquita1-1* (also called *cost1),* by growing mutant seedlings together with the wild type to ensure uniform soil water availability across genotypes. Our results demonstrate that the dwarf phenotype conferred by loss of CHIQ1 function results in constitutively lower water usage, but not increased drought resistance.

## Introduction

Among the various stresses plants endure in both natural and cultivated environments, drought stress has the greatest impact on plant productivity (Hu and Xiong, 2014). From an agricultural context, drought can be defined as the state of insufficient water availability to sustain maximum plant growth (Deikman et al., 2012). The impact of drought on global crop yields has intensified recently and is projected to intensify even more so in the future (Lesk et al., 2016; Leng and Hall, 2019). Identifying and engineering more drought resistant crops is therefore necessary to provide sufficient food to a growing population (Godfray et al., 2010).

Plants employ various mechanisms in response to drought. The specific responses to drought are influenced by the degree of stress, plant species, genotype within species, and developmental stage (Hu and Xiong, 2014). Some species and genotypes respond by hastening the completion of their life cycle before the onset of more severe stress (‘drought escape’) (Lawlor, 2013). Others respond by conserving or acquiring more water (‘drought avoidance’), or by maintaining metabolic homeostasis to prevent or repair damaged cells and tissues (‘drought tolerance’) (Lawlor, 2013). The many terms used throughout the literature to describe plant responses to water deficit (e.g. drought resistance, drought tolerance, drought avoidance) are often used interchangeably, resulting in ambiguity and a deviation from established terminology (Lawlor, 2013; Blum, 2014). This problem is compounded by results which could imply one or more forms of drought resistance (which encompasses escape, tolerance, and avoidance (Lawlor, 2013)) depending on the available data. For example, in response to reduced soil water availability, a plant could respond by increasing root growth (a drought avoidance response; (Levitt, 1980)) or via osmotic adjustment to maintain cell turgor (a drought tolerance response; (Morgan, 1984)). Without establishing which of these mechanisms is involved, we cannot ascertain which specific drought resistance response is responsible for an observed phenotype.

Despite the well-reasoned need to evaluate drought responses of mutant lines at equal levels of drought stress as controls (Lawlor, 2013; Turner, 2019), there are many claims of increased drought resistance that do not include this essential comparison (for example (Zhang et al., 2009; Bao et al., 2020; Hong et al., 2020)). In all such cases, mutant seedlings that survived longer or had greater rates of recovery after drought were not grown in the same pots with control plants nor was percent soil water content (SWC) kept uniform across genotypes. Therefore, the plants were evaluated at potentially unequal levels of drought stress. This situation is particularly problematic for plants that may use water at different rates, such as dwarf plants.

To demonstrate how controlling for SWC can affect the onset and degree of stress symptoms, and ultimately interpretation of results, we re-evaluated the drought phenotype of a dwarf Arabidopsis mutant recently implicated in drought tolerance. Bao and colleagues implicated *constitutively stressed 1 (COST1*) (originally characterized and named *CHIQ1* (Bossi et al., 2017)) in drought tolerance when grown in pots separate from the wild type (Bao et al., 2020). Here, we reassessed the drought phenotype associated with loss of *CHIQ1* function when *chiq1-1* seedlings were grown together with the wild type. Contrary to the previous report (Bao et al., 2020), we found that *chiq1-1* plants do not exhibit increased resistance to drought, despite constitutive lower water usage, compared to the wild type. Our study provides an easily reproducible, low-cost experimental approach to more effectively identify drought resistant genotypes, which will advance efforts to breed more resilient crops against climate change.

## Results

### *chiq1-1* mutant plants do not confer more drought resistance than the wild type

We evaluated *chiq1-1’s* water requirements and survival during drought to determine whether *CHIQ1* is involved in drought resistance or if *chiq1-1* plants simply use less water. When grown in pots with only a single genotype (either all wild type or all *chiq1-1), chiq1-1* plants survive longer during drought than wild type plants (Fig. 1B), consistent with the previous study (Bao et al., 2020). We next asked whether this phenotype was due to increased resistance to drought, or rather due to differences in the rate of water use between genotypes. We found that *chiq1-1* plants take up less water from the soil under both well-watered and drought conditions based on daily SWC levels (Fig. 1C-D). Reintroducing wild type *CHIQ1* into the mutant background complemented the water-use and survival phenotypes observed in the *chiq1-1* null mutant (Fig. 1B-D).

**Figure 1.**
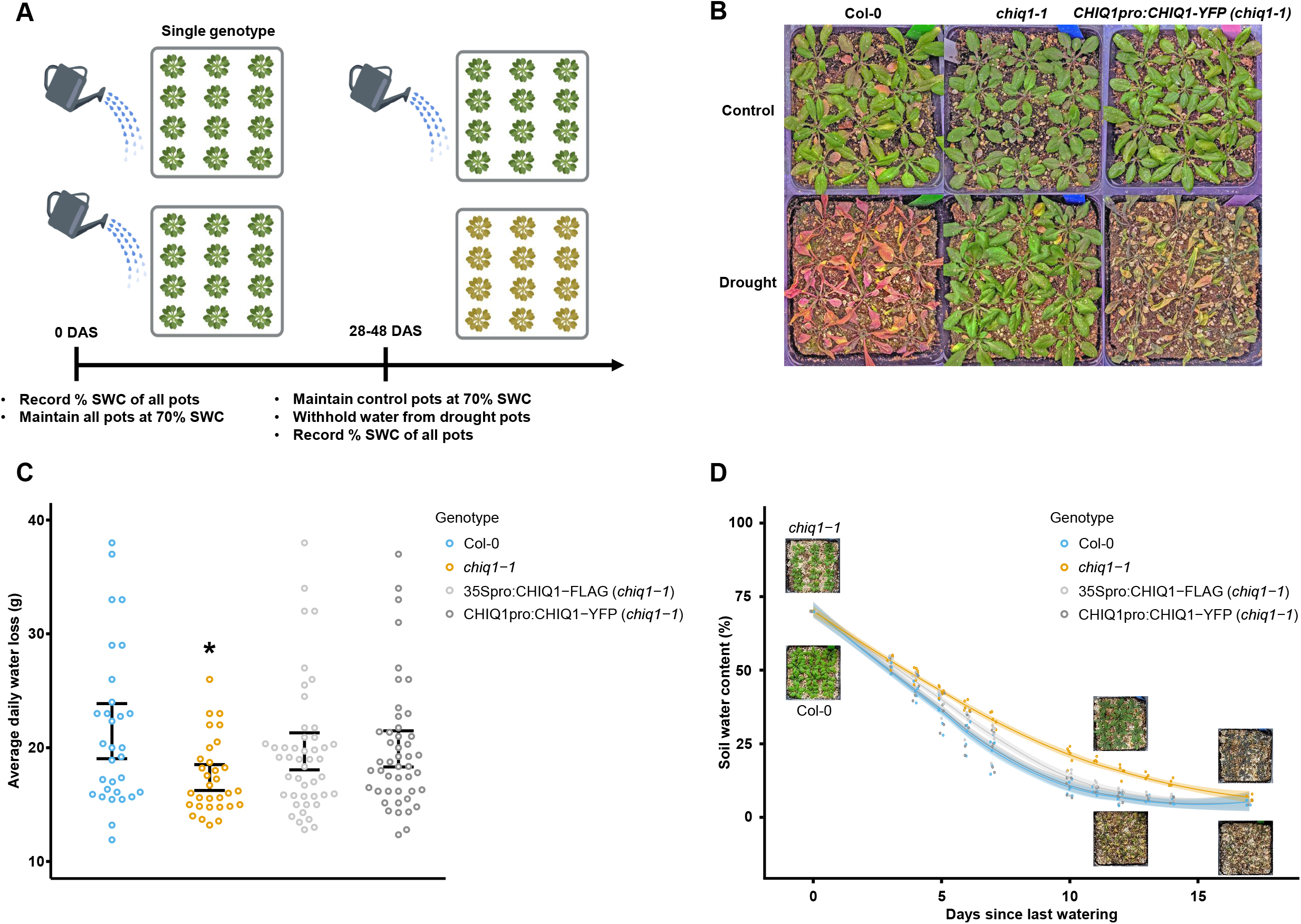
*chiq1-1* plants use less water and survive longer than the wild type when grown together. A) Timeline and description of experimental design in which each pot contained only a single genotype. B) Representative images of Col-0, *chiq1-1,* and the complemented line *proCHIQ1:CHIQ1-YFP* (in a *chiq1-1* background) grown in separate pots in control and drought conditions (12 days since last watering). C) Average daily water loss by genotype in well-watered (control) conditions (n = 31-46; N = 4). Black asterisk indicates statistical significance (p-value < 0.05) using Dunnett’s test with Col-0 as control. D) Percent soil water content by genotype during drought. Light-colored bands represent 95% confidence intervals of (n = 4-6; N = 2-3). Representative Col-0 and *chiq1-1* images are shown at 0, 12, and 17 days since the last watering. n = number of biological replicates per experiment; N = number of independent experiments.

When *chiq1-1* plants were grown in pots together with the wild type such that SWC was always equal for both genotypes, the visual onset of stress symptoms and duration of survival was uniform across genotypes (Fig. 2B, Movie 1). Photosystem II (PSII) quantum efficiency (F_v_/F_m_), a commonly used metric to quantify plant stress (Baker, 2008), and leaf relative water content (LRWC) as a percent of control decreased uniformly in both genotypes, when planted together, as a result of withholding water (Fig. 2C-D). Additionally, relative expression of the drought response genes RD22 and RD29A (Huang et al., 2018; Yu et al., 2019) was equal in both genotypes as a result of drought (Fig. 2E-F). Relative expression of COR15A, another commonly used drought response gene (Huang et al., 2018), was unchanged in either genotype in response to drought (Fig. S1). Together, these results indicate that *CHIQ1* is not involved in drought resistance, but rather that *chiq1-1* plants constitutively use less water, resulting in a slower decrease in soil water availability and a delayed onset of stress symptoms when grown separately from the wild type.

**Figure 2.**
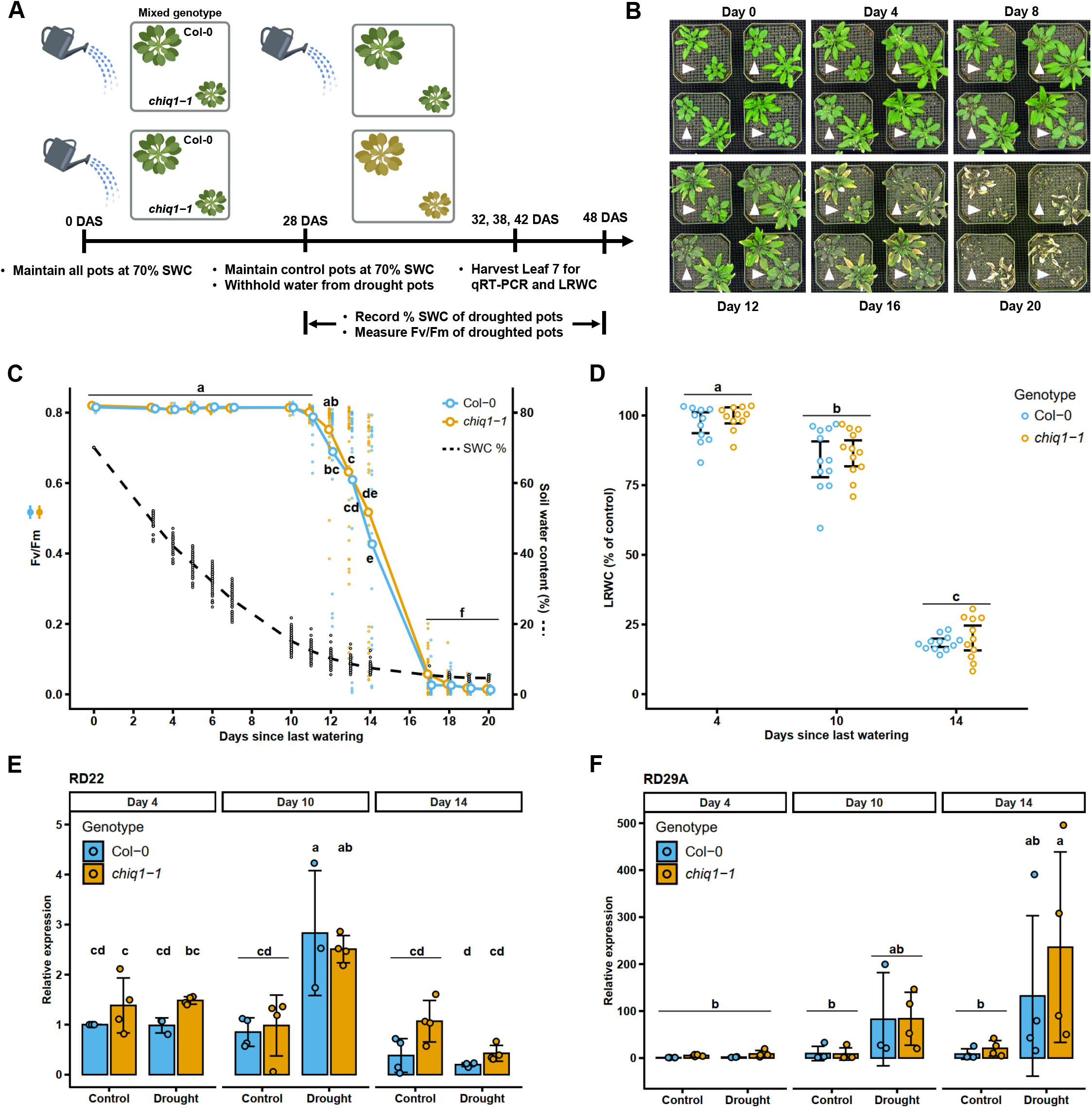
*chiq1-1* plants display equal drought resistance to the wild type when grown together. A) Timeline and description of experimental design in which each pot contained one Col-0 and one *chiq1-1* seedling. B) Representative images of pots containing one Col-0 and *chiq1-1* seedling over the course of drought. All 4 pots within each day panel are biological replicates. Day numbers represent days since last watering. Arrows point towards the *chiq1-1* seedling. C) Photosystem II quantum efficiency (F_V_/F_M_) as a function of drought (n = 14-46; N = 3-4). Soil water content % at each time-point is overlaid in the black dashed line (n = 32-48; N = 2-3). D) Leaf 7 relative water content as a percent of well-watered controls at 4, 10, and 14 days since last watering. Error bars represent the 95% confidence interval. E-F) Relative expression fold-change of desiccation marker genes RD22 and RD29A. Values were normalized by the housekeeping gene AT1G13320 and by Col-0; Day 4; Control using the ΔΔCt method. Each dot represents a pooled sample from an independent experimental repeat. Error bars represent the 95% confidence interval. Letters in C-F represent significantly different groups (p-value□<□0.05) as determined by two-(C,D) or three-way (E,F) ANOVA followed by Tukey’s HSD test. n = number of biological replicates per experiment. N = number of independent experiments.

## Discussion

Plants with reduced size often survive longer in response to water deprivation (Lawlor, 2013; Puértolas et al., 2017; Turner, 2019). We previously showed that *chiq1-1* plants have smaller organs than the wild type (Bossi et al., 2017; Bossi et al., 2021). In this study, we found that the reduction in plant size as a result of loss of *CHIQ1* function does not confer drought resistance. This is contrary to what was recently published (Bao et al., 2020), where wild type and *chiq1-1* plants were grown and droughted in different pots with the implicit assumption that SWC was equal in all pots after withholding water. This assumption can dramatically alter the conclusions drawn regarding drought resistance, as illustrated in this study. We showed that *chiq1-1* plants use less water than the wild type and therefore the SWC in pots containing only Col-0 or only *chiq1-1* was different as a function of time after withholding water. When we grew *chiq1-1* plants in the same pot as the wild type, such that both genotypes were always forced to cope with equal levels of SWC, *chiq1-1* plants were qualitatively and quantitatively no more resistant than the wild type to drought stress. The absence of a *chiq1-1* drought phenotype when grown together with the wild type was observed both physiologically and molecularly, the latter in direct contrast to the expression profiles of RD22, RD29A, and COR15A in the wild type and *chiq1-1* when grown and droughted in separate pots (Bao et al., 2020). This is not to say that *chiq1-1* is not potentially advantageous in an agronomic context (for example in a monoculture environment in which all plants are *chiq1-1).* Indeed, daily water usage in both well-watered and drought conditions demonstrates that the dwarf *chiq1-1* plants constitutively use less water than the wild type. However, when situated in an environment more competitive for water use, *chiq1-1* plants fare no better than their wild type neighbors. Our work highlights the importance of ensuring that comparisons between genotypes are made at equal levels of drought stress by subjecting all genotypes to uniform levels of stress. Drought is the single biggest source of crop production loss, resulting in billions of dollars in losses worldwide (FAO, 2021). The easily-reproducible and cost-effective methods we used to evaluate *chiq1-1’s* drought phenotype will serve as a valuable resource to the plant science community in designing future experiments to screen for genotypes more resilient to drought.

## Materials and Methods

### Plant materials and growth conditions

Water content of fresh PRO-MIX HP Mycorrhizae potting soil (Premier Tech Horticulture, Quakertown, PA) was determined by drying 3 samples of fresh soil at 45°C for 1 week. Average water content of fresh soil was calculated as dry weight/fresh weight. To determine soil water holding capacity (100% SWC), 8 pots were filled with fresh soil, weighed, saturated with water, covered, and then left to drip until pots reached pot capacity (cessation of dripping). They were then weighed again to determine the average water-holding capacity of the soil.

Wild type *Arabidopsis thaliana* accession Columbia-0 (Col-0) and *chiq1-1* mutant (SALK_064001) seeds were obtained from the Arabidopsis Biological Resource Center (ABRC). *CHIQ1* complementation lines were obtained as described in (Bossi et al., 2021). Seeds were stratified in water at 4°C for 4 days before planting. Seedlings were grown in a growth chamber under a 16:8 hour light:dark cycle at 22°C, 40% RH, and ~100 μmol m-^2^ s-^1^ photosynthetic photon flux density (PPFD) measured at pot-level. Flats were rotated daily Monday-Friday to avoid positional effects. All experiments were conducted at least three times independently.

### Single genotype per pot drought experiment

All pots were filled with an equal amount of PRO-MIX HP Mycorrhizae potting soil by weight. Seeds were planted such that each pot contained 12 seedlings of a single genotype (Col-0, *chiq1-1, proCHIQ1:CHIQ1-YFP* (in a *chiq1-1* background), or *35Spro:CHIQ1-FLAG* (in a *chiq1-1* background). To obtain 12 seedlings per pot for the single-genotype per pot experiments, 3-4 seeds were planted in each of 12 locations within a pot. After seeding, pots were put into flats and were covered for 1 week, after which covers were removed and each pot was thinned to contain 12 seedlings. At 28 days after sowing (DAS), pots were either subjected to drought (total withholding of water) or were maintained at 70% SWC as controls. All pots were weighed daily Monday-Friday to determine water loss in both control and drought conditions. Statistical differences in weekday water usage per day across genotypes was determined by one-way analysis of variance (ANOVA) followed by Dunnett’s test (P□<□0.005) setting Col-0 as control and using the DunnettTest() function within the DescTools package in R version 3.6.3.

### Multiple genotypes per pot drought experiment

For experiments directly comparing drought resistance between Col-0 and *chiq1-1* plants, one seedling each of Col-0 and *chiq1-1* were planted together in individual pots. All pots were filled with an equal amount of PRO-MIX HP Mycorrhizae potting soil by weight. At 28 DAS, half of the pots were subjected to drought and the other half were maintained at 70% SWC as controls. Droughted pots were weighed daily Monday-Friday to determine % SWC as a function of time.

### Image capture and timelapse generation

Images were taken every 2 hours from directly above pots using a Raspberry Pi Zero W (Raspberry Pi Foundation, Cambridge UK) and an Arducam M12 lens (model B0031; https://arducam.com). Images were captured using the camera.capture() Python function and were taken at 2-hour intervals using the command-line job scheduler, crontab (Unix). To remove lens distortion, images were corrected in Adobe Photoshop CS6 (Adobe Systems, Inc., San Jose, CA, USA) using the “Lens correction” feature. All images were then stitched together into a time series video using Davinci Resolve 17 (Blackmagic Design, Port Melbourne, Victoria, Australia).

### Chlorophyll fluorescence measurements

Chlorophyll fluorescence parameters were measured between 9:30-10:00am on the 7th true leaf of each sample using a chlorophyll fluorometer (OS30p+, Opti-Sciences, Inc. Hudson, New Hampshire). After dark-adapting leaves for 30 minutes, a weak modulated light (0.1 μmol m^-2^ s^-1^ PPFD) was applied to measure minimum fluorescence (F_0_). Maximum fluorescence (F_M_) was measured after applying a saturating light pulse (6000 μmol m^-2^ s^-1^ PPFD) of 1 second to the sampled region. Photosystem II quantum efficiency (F_V_/F_M_) was calculated as (F_M_ - F_0_)/F_M_. Statistical differences in F_V_/F_M_ values between genotypes as a function of time were determined by two-way ANOVA followed by Tukey’s honestly significant difference test (p□<□0.05) using the lsmeans() function within the lsmeans package in R version 3.6.3.

### qRT-PCR

At 4, 10, and 14 days of drought treatment, a 0.38 cm^2^ disc was excised from the 7th true leaf of well-watered and droughted plants and immediately frozen in liquid nitrogen. Leaves from 4 plants of the same genotype, condition, and time point were pooled for each experiment. Four independent experiments were conducted. Frozen tissue was homogenized and mRNA was extracted using the QIAGEN Plant RNeasy Plant Mini Kit (QIAGEN, Hilden, Germany). mRNA was reverse transcribed using the Protoscript II First Strand cDNA Synthesis Kit (New England BioLabs, Ipswich, MA, USA). qRT-PCR reactions were performed in LightCycler 480 96-well plates (Roche Diagnostics, Indianapolis, IN, USA) using a 10 minute pre-incubation period at 95°C followed by 40 cycles of denaturation for 5 seconds at 95°C, annealing for 10 seconds at 60°C, and extension for 10 seconds at 72°C. Amplification of target genes was measured each cycle by SYBR Green I fluorescent dye. All qRT-PCR reactions were performed in a Roche LightCycler 480 II (Roche Sequencing and Life Sciences, Indianapolis, IN, USA) using the LightCycler 480 SYBR Green I Master reaction mix. Transcript abundance was normalized by the housekeeping gene AT1G13320 (Czechowski et al., 2005) and relative expression fold-change was calculated using the delta delta Ct (ΔΔCt) method (Livak and Schmittgen, 2001). Statistical differences in relative expression fold-change were determined by three-way ANOVA followed by Tukey’s honestly significant difference test (P□<□0.05) using the lsmeans() function within the lsmeans package in R version 3.6.3. All primers are listed in Table S1.

### Leaf relative water content

The remaining leaf tissue from which discs were cut out for qRT-PCR was weighed, floated on deionized water for 2 hours, reweighed, and then dried for 48 hours at 65°C. LRWC was calculated as (pre-float weight - dry weight) / (post-float weight - dry weight) * 100. Statistical differences in LRWC as a percent of control were determined by two-way ANOVA followed by Tukey’s honestly significant difference test (P□<□0.05) using the lsmeans() function within the lsmeans package in R version 3.6.3.

## Acknowledgments

We thank the Arabidopsis Biological Resource Center (ABRC) for providing *chiq1-1* (SALK_064001) mutant seeds, A. Malkovskiy for helpful advice on imaging, Y. Dorone for helpful suggestions, and I. Villa and G. Materassi-Shultz for plant growth facility support. This work was done on the ancestral land of the Muwekma Ohlone Tribe, which was and continues to be of great importance to the Ohlone people.

## Authors’ contributions

F.B., S.Y.R., and D.G. conceived the project. D.G. performed the drought studies. F.B. and S.Y.R. provided intellectual contribution and input into manuscript organization. D.G. wrote the manuscript and F.B. and S.Y.R. edited the manuscript.

## Competing interests

Authors declare no conflict of interest.

## Funding

This work was supported by grants from the National Science Foundation (IOS-1546838, IOS-1026003) and the US Department of Energy, Office of Science, Office of Biological and Environmental Research, Genomic Science Program grant nos. DE-SC0018277, DE-SC0008769, DE-SC0020366, and DE-SC0021286.

## Data availability

All study data are included in the main text and supporting information.

## Authors’ information

Correspondence and requests for materials should be addressed to S.Y.R.

---

Movie 1

**Visual onset and progression of drought stress is uniform between the wild type and *chiq1-1* plants when grown in shared pots.** Timelapse videos of pots containing one Col-0 and *chiq1-1* seedling. Orange arrows point towards the *chiq1-1* seedling. All pots are biological replicates. Video begins on the last day of watering (28 DAS) and ends 20 days later. Three independent experiments were conducted.

**Supplementary Information**

**Figure S1**

Relative expression fold-change of desiccation marker gene COR15A. Values were normalized by the housekeeping gene AT1G13320 and by Col-0; Day 4; Control using the ΔΔCt method. Each dot represents a pooled sample from an independent experimental repeat. Error bars represent the 95% confidence interval. Statistical significance was determined by three-way ANOVA followed by Tukey’s HSD test.

**Table S1.**
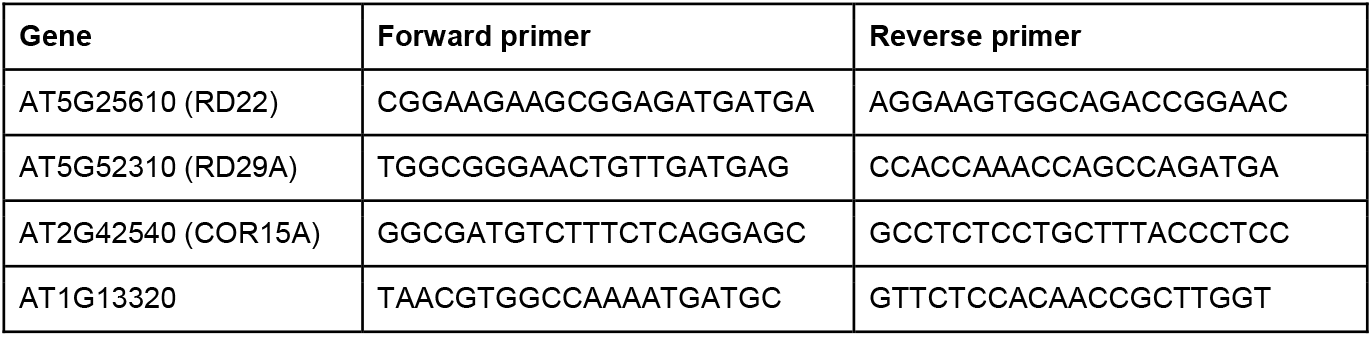
Primers used for qRT-PCR.

## Notes

### Competing Interest Statement

The authors have declared no competing interest.

### Summary of Updates

Inclusion of leaf relative water content and qRT-PCR data

