## Supplementary figures and images for "Uncoupling differential water usage from drought resistance in a dwarf Arabidopsis mutant"

### Supplemental Figure 1

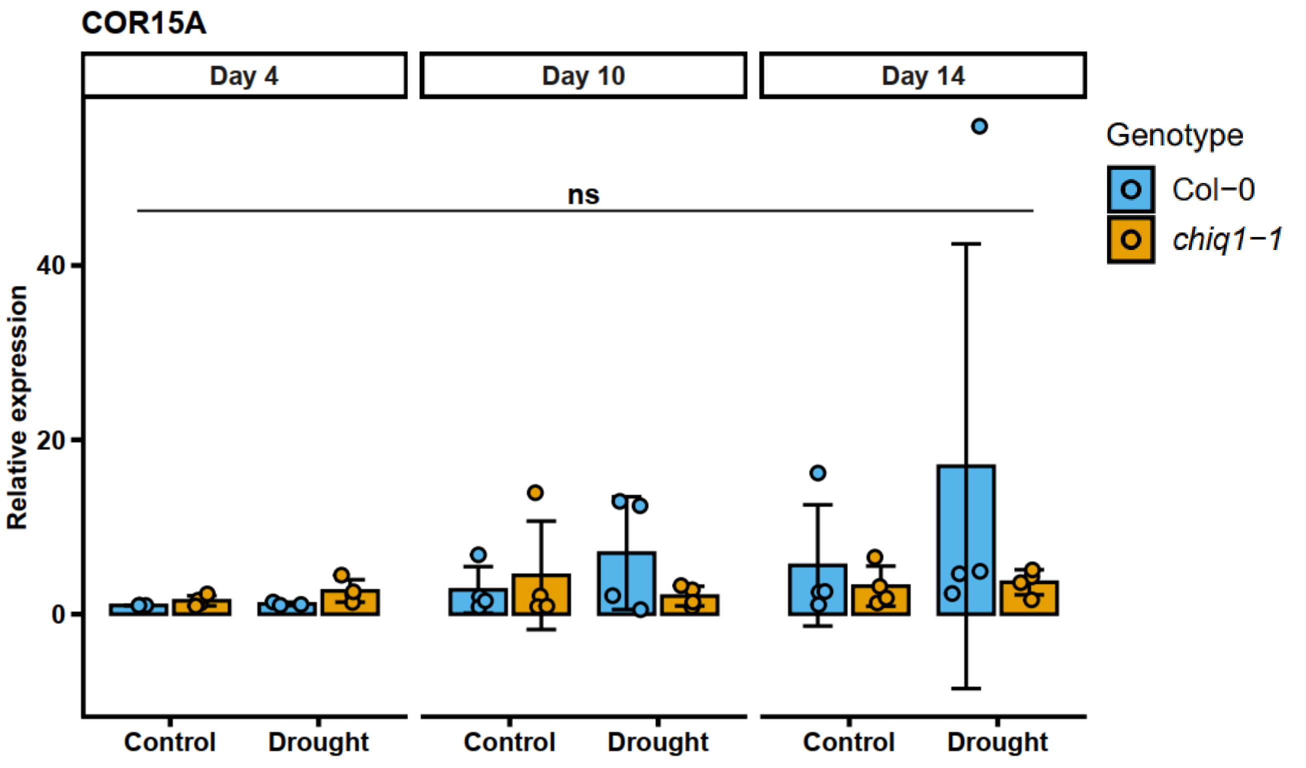
